# Targeted Neuromodulation of Perceptual Decision-Making Networks Causally Dissociates Sensory and Metacognitive performance

**DOI:** 10.1101/2025.05.15.653831

**Authors:** Paolo Di Luzio, Luca Tarasi, Alessio Avenanti, Juha Silvanto, Alejandra Sel, Vincenzo Romei

**Affiliations:** Centre for Brain Science, Department of Psychology, University of Essex, Wivenhoe Park, Colchester CO4 3SQ, UK; Centro studi e ricerche in Neuroscienze Cognitive, Dipartimento di Psicologia “Renzo Canestrari”, Alma Mater Studiorum Università di Bologna, Campus di Cesena, Viale Rasi e Spinelli 176, 47521 Cesena, Italy; Essex ESNEFT Psychological Research Unit for Behaviour, Health and Wellbeing, University of Essex, Wivenhoe Park, Colchester, CO4 3SQ, UK; Centro de Investigación en Neuropsicología y Neurociencias Cognitivas, Universidad Católica del Maule, 3460000 Talca, Chile; Center for Cognitive Brain Sciences, University of Macau, Avenida de Universidade, Macau, China SAR; Facultad de Lenguas y Educación, Universidad Antonio de Nebrija, Madrid 28015, Spain

## Abstract

Perceptual judgements and their subjective evaluation (i.e., metacognition) are considered tightly coupled aspects of decision-making. Yet, emerging evidence suggests that distinct neural mechanisms underlie these processes. In motion perception, studies have identified a neural network involving visual and associative parietal areas supporting these abilities. Whilst early (V1/V2) and specialized (V5/MT+) visual areas are associated with motion discrimination accuracy, the intraparietal sulcus (IPS/LIP) plays a critical role in the formation of subjective confidence for sensory decision, given its functionally coupling with the V5/MT+–V1/V2 network. Previous studies have consistently reported that increasing connectivity in the human V5/MT+-to-V1/V2 pathway by means of cortico-cortical paired associative stimulation (ccPAS) – a neurostimulation protocol inducing Hebbian-like changes in neuronal pathways – improves our perceptual ability to discriminate motion. Notably, we recently demonstrated that strengthening V5/MT+ influence over V1/V2 also leads to overconfidence – altering metacognitive bias for motion discrimination, suggesting that the manipulation of V5/MT+–V1/V2 network might functionally affect IPS/LIP activity. Here, we directly investigated this possibility by first strengthening the V5/MT+-V1 pathway with ccPAS, and subsequently interfering with IPS/LIP activity by means of continuous theta-burst stimulation (cTBS). In line with previous evidence, our results corroborate that enhancing V5/MT+-V1/V2 connectivity affects both motion discrimination and metacognitive bias. Crucially, IPS/LIP disruption through cTBS selectively affects metacognitive performance while leaving motion discrimination unaltered. These findings highlight distinct neural substrates for perceptual sensitivity and visual metacognition linked to V5/MT+ and IPS/LIP, respectively, opening to potential tailored applications of non-invasive brain stimulation (NIBS) to functionally improve sensory and metacognitive decision-making processes.

## Introduction

Humans tend to show proficiency in estimating the physical properties of a stimulus, such as motion direction, and consistently monitor the accuracy of their decisions. However, the accuracy of perceptual judgments and the corresponding level of confidence (i.e., metacognition) do not always align [1]. This misalignment may be attributed to the fact that sensory accuracy and metacognitive abilities are mediated by distinct brain regions within the perceptual system [2,3]. This is consistent with a neural architecture where choice and confidence emerge from parallel computations, as predicted by dual-channel models [4,5]. Accordingly, behavioural dissociations between visual sensitivity and metacognitive indices are often reported [6–8], demonstrating how these two dimensions of decision-making can be independently examined [9–11].

In animals, perceptual decision-making has been extensively investigated in motion discrimination, showing that middle temporal (MT+) and lateral intraparietal (LIP) are fundamental areas involved in perceptual judgements [12,13]. Specifically, the impairment of neurons in MT+ of non-human primates selectively hinders sensory performance [13,14], while inhibiting activity in the LIP – whose neurons integrate signals supporting decision confidence [12] – has no impact on perceptual sensitivity [13]. These findings strongly suggest that MT+ and the LIP play distinctive functional roles in decision-making.

Consistently, recent evidence from our group suggests a partial dissociation between sensory and metacognitive decision systems in humans [15], highlighting the functional role of the network connecting the extrastriate (V5/MT+), and the intraparietal sulcus (IPS) – the human homologue of LIP [16,17] – with early visual cortices (V1/V2). Specifically, selectively strengthening the V5/MT+-to-V1/V2 and the IPS/LIP-to-V1/V2 pathways with cortico-cortical paired associative stimulation (ccPAS) –a neurostimulation protocol designed to induce Hebbian-like changes in neuronal pathways [for reviews 18–20]– distinctively affects the perceptual behaviour of the participants. Enhancing V5/MT+ influence over V1/V2 with ccPAS results in reduced motion perception threshold (i.e., finer sensory discrimination [21,22]) and a generalised increase in confidence levels (i.e., changes in metacognitive bias) [15]. By contrast, increasing synaptic plasticity in the route connecting the IPS/LIP to V1/V2 with ccPAS, leads to a selective modulation of metacognitive efficiency (i.e., level of confidence matching the sensory accuracy), while leaving sensory performance per se unaltered [15]. Increases in motion perception and changes in metacognitive bias resulting from enhanced functional influence of V5/MT+ over V1/V2 have been later observed by another independent research group [23]. This study also recorded electrophysiological (EEG) responses [23] showing that changes in metacognitive bias (i.e., overconfidence) following ccPAS_V5/MT+--V1/V2_ are accompanied by enhanced bottom-up signals from V1/V2-to-IPS/LIP. These observations aling with previous functional imaging findings of the ccPAS effect on different networks (e.g., motor-premotor), showing that the increased interregional connectivity following the stimulation is prominent between the stimulated areas themselves, but also extends to other closely interconnected cortical areas [24].

It is thus possible that the perceptual overconfidence reported when manipulating V5/MT+--V1/V2 connectivity might be partially driven by increases in the functional influence that this network exerts over IPS/LIP [23], as allowed by their anatomical connections [25]. Accordingly, microstimulation of monkeys MT+ during motion discrimination is able to affect their sensory performance and confidence [26], suggesting that increased perception may secondarily influence choice certainty [27]. This possibility seems particularly plausible in light of evidence that perceptual decisions are the result of functional interactions encompassing a broad system of regions, with a hierarchical elaboration of sensory signals [28–30]. We therefore hypothesized that enhancing interregional connectivity from V5/MT+ to V1/V2 would increase its functional influence over IPS/LIP, thus affecting metacognitive bias. Crucially, after strengthening V5/MT+-to-V1/V2 connectivity with ccPAS, we disrupted IPS/LIP activity to test its specific contribution to perceptual metacognitive abilities. Since IPS/LIP is not expected to be directly involved in motion sensitivity, its disruption should not affect basic perceptual performance. Here, using a multimodal, non-invasive brain stimulation approach, we demonstrate the functional independence of human neural circuits involved in perceptual sensitivity and metacognition.

## Methods

### Experimental paradigm

We applied a repeated dual coil paired stimulation over V5/MT+ and V1/V2 with the aim of strengthening their connectivity (ccPAS_V5/MT+--V1/V2_, see *ccPAS*) in two groups of participants (see *participants*). To assess the impact of this intervention on motion perception and visual metacognition, participants completed a motion discrimination task (see *Behavioural task*) in two initial blocks: before (Baseline) and 30 minutes (T30) after offline ccPAS. Increases in visual motion perception and metacognition after strengthening the V5/MT+-to-V1/V2 pathway with ccPAS are typically peaking at 30 min after ccPAS [18]. Having induced perceptual and metacognitive alterations with ccPAS_V5/MT+--V1/V2_ [15,21,23], we then applied continuous theta-burst stimulation (cTBS, see *Continuous theta-burst stimulation*) over IPS/LIP with the aim of interfering its activity in a subgroup of participants (cTBS_Active_, N=18). By contrast, sham cTBS over IPS/LIP was administered to a distinct group of participants (cTBS_Sham_, N=18). If the altered metacognition observed following ccPAS_V5/MT+--V1/V2_ results from enhanced functional influence of V1/V2 over IPS/LIP [23], then disrupting IPS/LIP activity with cTBS should selectively impair metacognitive abilities without affecting discrimination performance. Conversely, given the expected duration of ccPAS effects [18,21], both perceptual and metacognitive alterations should persist after sham IPS/LIP inhibition. Specifically, we reassessed performances in both groups ~5 minutes after active or sham cTBS in a third test block (T60; Fig.1A). Our multimodal non-invasive brain stimulation approach provides the ideal testbed for our hypotheses. In particular, this allowed us to investigate the functional dissociation between the effective modulation of V5/MT+ on V1/V2, and its operational influence over the IPS/LIP region in sensory and metacognitive performances, respectively. We assessed motion threshold and confidence levels (i.e., metacognition) focusing on the relative changes observed in the blocks at 30 and 60 minutes after ccPAS, compared to baseline (see *Statistical analysis*).

**Figure 1.**
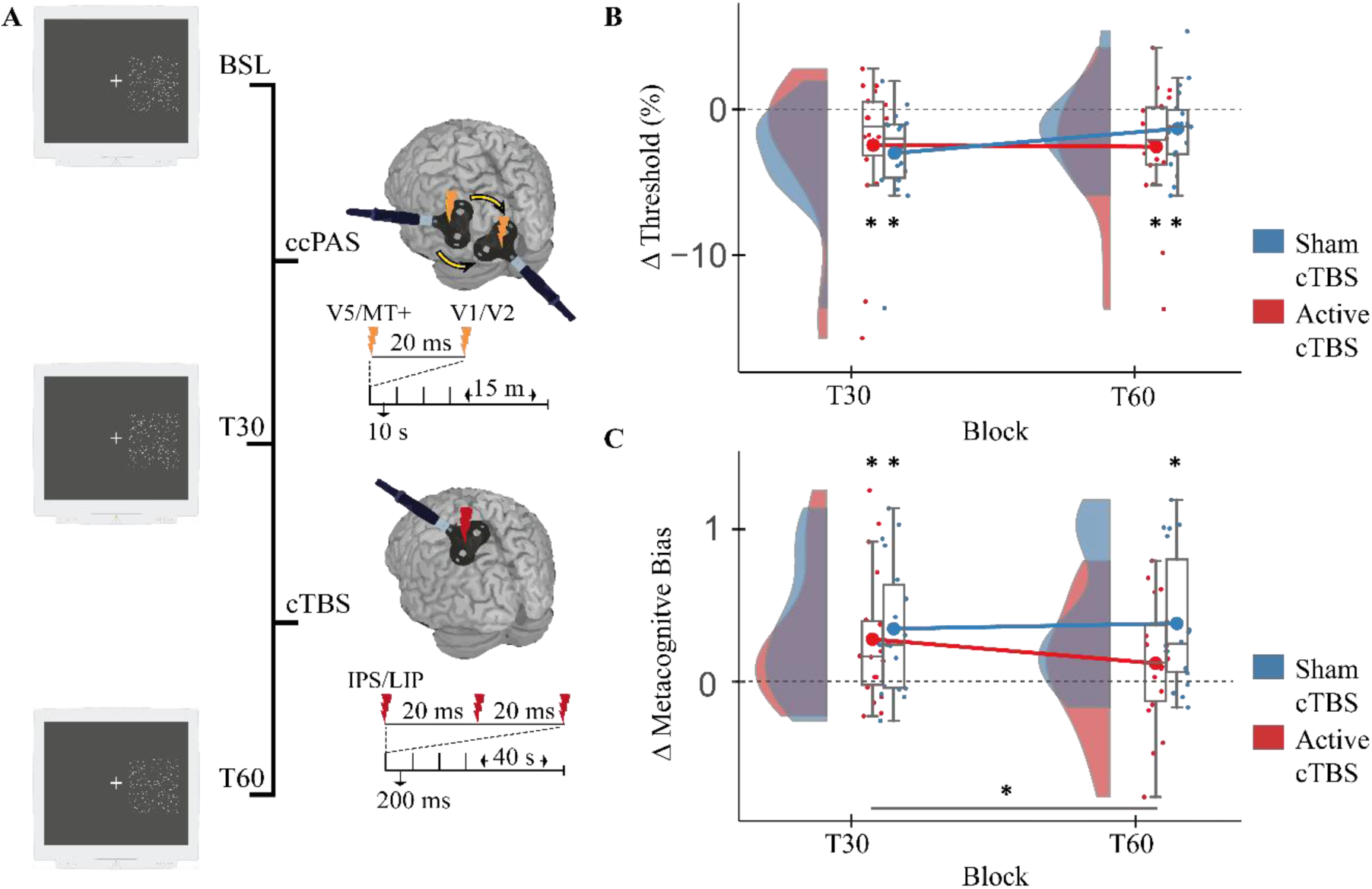
(**A**) Representation of the experimental procedure: participants (N= 36) performed the motion discrimination task prior to (BSL) and 30 minutes after (T30) the V5-V1 ccPAS protocol. One group of participants (N=18) underwent a cTBS protocol (Active_cTBS) targeting IPS/LIP and one group a sham protocol (Sham_cTBS), after which they were re-tested on the motion discrimination task 60 minutes from the ccPAS administration (T60). **(B)** Changes in motion sensitivity threshold following ccPAS and cTBS. Half-density and Box-plots with individual data points represent raw, mean and median of delta perceptual threshold at T30 and T60 for active (red) and sham (blue) condition of cTBS; Asterisks show corrected one-sample t-test against null values (p<0.05). **(C)** Changes in metacognitive bias following ccPAS and cTBS. Half-density and Box-plots with individual data points represent raw, mean and median of delta perceptual confidence at T30 and T60 for active (red) and sham (blue) condition of cTBS; Asterisks show significant (p<0.05) one-sample t-test and post-hoc comparison.

### Participants

Thirty-six right-handed healthy volunteers (17 men; mean age ± SD = 23.47 ± 5.21) took part in the study. Sample size was determined according to our previous report [15], aiming for a similar number of participants given the between-group design. Participants were evenly assigned to an experimental and a control condition, *Active_cTBS* and *Sham_cTBS*, respectively, and were matched for numerosity and gender. None of them had previously reported mental or neurological pathologies or contraindications to the TMS administration procedure [31]. Each participant provided their informed consent prior to the study and all procedures received the approval of the local bioethics committee of the Department of Psychology, University of Bologna.

### Behavioural task

The experimental task, a modified version of the random-dot kinematogram [15], was projected on a 18-inch CRT monitor (with a resolution of 1280 x 1024 pixels and a refresh rate equal to 85 Hz) distant ̴ 57 cm from the participant. The stimuli were generated and presented using the MATLAB statistical analysis environment (version 2019a, The MathWorks Inc., Natick, MA) and the Psychophysics Toolbox [32]. At the beginning of each trial, a fixation cross in the centre of the screen was displayed for 500 ms; followed by a matrix of moving dots plotted within a square region of 12.8 x 12.8 visual degrees (°) to the right of the fixation point, lasting 400 ms. A fixation cross was then displayed again until the manual response, and a subsequent screen showing a confidence rating was presented at the end of the trial. The motion stimulus consisted of 400 white dots (each dot being 6 pixels in size) that could move leftwards or rightwards within a range of 10 different percentages of coherence (0, 2, 4, 6, 12, 16, 20, 35, 50, 80). The coherence of motion reflected the proportion of dots representing the signal (e.g., 20%, 80 dots heading left or right, 320 randomly moving). During task presentation, participants were asked to keep their gaze on the fixation cross and, at stimulus offset, they were asked to decide which motion direction they perceived, choosing between two alternative responses (left-right) by pressing the corresponding arrow on a keyboard; these judgments were used to fit a psychophysical curve in order to measure each participant performance accuracy. Participants were then asked to rate, on a discrete four-point scale, their level of confidence on the previously expressed judgment (1: totally unconfident, 2: unconfident, 3: quite confident, 4: totally confident) by selecting the corresponding number keys; this measure was used to compute metacognitive indices (*see Statistical analysis*). Each block included 20 repetitions for each of the 10 coherence percentages in the two possible motion directions (leftwards or rightwards), for a total of 400 trials over a period of approximately 12 minutes.

### ccPAS protocol

The ccPAS procedure required the use of a pair of figure-of-eights iron-branding coils (50-mm) connected to a Magstim BiStim2 stimulator assembled from two monophasic 200 modules. The application of the protocol took about 15 minutes. Ninety pairs of pulses were applied at a constant frequency of 0.1 Hz, to minimize temporal summation effects [33]; each pair consisted of two monophasic transcranial magnetic stimulation (TMS) pulses, automatically triggered by a MATLAB interface. Both experimental groups underwent the same ccPAS protocol. The areas of interest for the stimulation were V5/MT + of the left hemisphere and bilateral V1/V2. The coils were oriented tangentially to the scalp with a stimulation intensity equal to 60% of the maximum stimulator output (MSO), in line with previous studies using fixed intensity for visual cortices [34–37]. In order to target V5/MT +, the coil was positioned 3 cm dorsally and 5 cm laterally to the inion, corresponding to the scalp position where, on average, TMS is able to induce moving phosphenes [38] and modulate visual motion perception [39]. In order to stimulate V1/V2, the coil was positioned 2 cm dorsal to the inion, corresponding to the area of the scalp whose stimulation typically elicits phosphenes. The temporal order of TMS pulses involved V5/MT+ stimulation first followed by V1/V2 stimulation. The inter-pulse interval (IPI) between pulses was set to 20 ms, in order to elicit consequential pre- and post-synaptic activation of the cortical areas under examination, based on the physiological latency of these reentrant projections [22,38].

### Continuous theta-burst stimulation (cTBS)

Sixty minutes after the administration of the ccPAS protocol, the participants underwent the cTBS session according to the condition they were assigned. For the *Active_cTBS* condition, stimulation was applied to modulate the activity of IPS in an inhibitory fashion, using a Magstim Rapid2 biphasic stimulator connected to a butterfly-shaped coil (70 mm). The stimulation train consisted of 600 pulses (cTBS-600) administered for ~45 seconds [40]. Within the train, bursts were repeated at 200 ms intervals and each of them consisted of three pulses released at a frequency of 50 Hz. In line with the literature, the stimulation intensity was imposed to be subthreshold, 80% of the phosphenes perception threshold [41–44]. Assuming an average PT of ~60% MSO [e.g., 45], this was consequently adjusted to a conservative value of 45% (MSO), which is consistent with reported intensities for cTBS of the visual cortex [42–44,46]. Coil was held tangential to the scalp with an anterior-to-posterior orientation. The inhibitory cTBS protocol has been shown to be able to exert a long-term depressive effect (LTD) of the cortical area subjected to stimulation for a period of more than 30 minutes [47]. The targeted site, corresponding to the coordinates P3 (international system 10-20), coincided with the IPS/LIP areas of the left hemisphere with a variability of less than 2 cm [48,49], thus allowing to minimize the margin of error between subjects. For the *Sham_cTBS*, all the stimulation parameters were the same as for the active condition, except for the coil surface that was tilted perpendicularly to the scalp, in order to avoid any magnetic discharge to be delivered on the cortical target. A neuronavigation software (SofTaxic, E.M.S., Bologna, Italy) combined with a 3D optical digitizer (Polaris Vicra, NDI, Waterloo, Canada) was used to control for the consistency of TMS pulse delivery over the marked scalp position throughout the experiment and to store average MNI coordinates of the cortical site. The SofTaxic software estimated the volume of magnetic resonance images of the participant’s head by means of a warping procedure, on the basis of the participant’s skull landmarks (nasion, inion, and two preauricular points) and a set of 65 points providing a uniform representation of the scalp.

### Statistical analysis

The collected data were plotted on a Cartesian plane in which the abscissa axis represented the coherence level of the movement (0–80%) and the ordinate axis represented the percentage of performance accuracy. The distribution of the variable simulated a psychophysical curve of sigmoid shape. Between the accuracy values of 50% (corresponding to a totally random coherence of motion) and 100% (corresponding to 80% coherence of motion) the curve tended to grow, as predicted based on the results of previous research [39]. The data were fitted through a non-linear function modeled on the logistic curve:

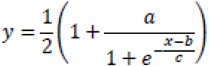

in which “a” represents the value of the upper horizontal asymptote; “b” represents the value at which the function changes its trend (inflection point), halfway between the lower and upper asymptotes; “c” represents the value that describes the direction and steepness of the curve (slope). For each participant in each Block, the value of the inflection point was calculated, corresponding to the intersection between the accuracy value of 75% and the relative level of coherence of the movement. As a result, the motion sensitivity threshold was determined, the value of which corresponded to the minimum percentage of coherent movement necessary to perform the motion discrimination task with an accuracy of 75%. In order to investigate the modulation of visual metacognition throughout the experimental Blocks, depending on the stimulation protocol, we evaluated different indices of metacognition. Specifically, by considering the level of performance in trials encompassing the individual motion threshold we obtained scores for metacognitive bias (i.e., average confidence level), metacognitive sensitivity (i.e., meta-d’, the sensitivity of confidence ratings), and efficiency (i.e., meta-d’-d’, how much the subjective sensitivity differs from the perceptual sensitivity) [50,51]. We computed the metacognitive bias of each subject as their average confidence, the higher the value the more confident the participant. Confidence ratings were then fitted through a single-subject Bayesian estimation approach [50] to estimate second-order (metacognitive) sensitivity expected to occur given the observed first-order (i.e., motion sensitivity) performance. This represents a signal detection theory (SDT) approach [51] to obtain indices of metacognitive sensitivity (i.e., how much sensory information participants can use to give confidence) and metacognitive efficiency (defined as difference between meta-d’ and d’), which normalizes metacognitive sensitivity (meta-d’) by task performance (d’) and provides the participants ability to calibrate confidence given their performance. Higher values of sensitivity means that confidence ratings are more informative regarding the accuracy of one’s judgments; conversely, the closer to zero the higher the metacognitive efficiency. The Bayesian estimation we adopted is more robust to variability in trial numbers and does not apply correction for missing cases relatively to previous implementation of meta-d’[50].

All the analyses were performed using SPSS 23 (IBM) statistical software and Matlab v2019b. We systematically tested group differences using normalized values (baseline-corrected, POST-PRE) by means of repeated measures ANOVAs considering the within-subject factor Block (Block_T30_; BlockT60) and between-subject factor cTBS (cTBS_Active_; cTBS_Sham_). For all the indices considered, performances were preliminary evaluated to assess the group’s homogeneity at baseline (Fs<.4, ps>.53). Control analysis using raw pre- and post-stimulation performance were performed considering factors of Block (Block_BSL_, Block_T30_, Block_T60_) and cTBS (Active, Sham).

Post-hoc test with Duncan [52] method were performed for between and within group comparisons following ANOVAs; Bonferroni-Holm corrected one-sample t-tests were used to contrast selected means against null values.

## Results

Firstly, the assessment of motion perception performance, conducted on normalized value of sensitivity, revealed the absence of interaction between the two neurostimulation protocols or effects across the blocks (Fs<4.03; ps>0.053; Fig.1B). This tendency towards significance motivated further analyses on raw motion discrimination to disambiguate the effect of ccPAS and cTBS on performance, revealing the expected reduction of sensory thresholds (Main effect of Block: F(2,68)=11.21; p < .001; ηp^2^=.25, Fig.2C). Compared to baseline (mean= 12.5±1.5) the decrease of thresholds emerged in Block_T30_ (mean= 8.43±0.9, p<.001), it persisted in Block_T60_ (mean= 9.6±1.3, p<.01) and was comparable between these two blocks (p=.19). The sensory improvement was present independently of whether active (T30= −2.48±1.3; T60= −2.73±1.1) or sham cTBS (T30= −2.98±0.82, T60= −1.35±0.6) were administered (Fs<1.21; ps >.30, Fig 2A-B), and thus not being affected by IPS inhibition. This proved that the stimulation functional manipulation of V5/MT+-to-V1/V2 reentrant network is effective in improving motion sensitivity up to 60 minutes, denoting only a non-significant attenuation of efficacy according to the time elapsed from ccPAS (Fig 2C), in line with previous reports showing maximum effects at 30 minutes from stimulation [16]. As hypothesized, this also confirms that IPS/LIP perturbation is insufficient to produce significant alterations in motion sensitivity [15].

**Figure 2.**
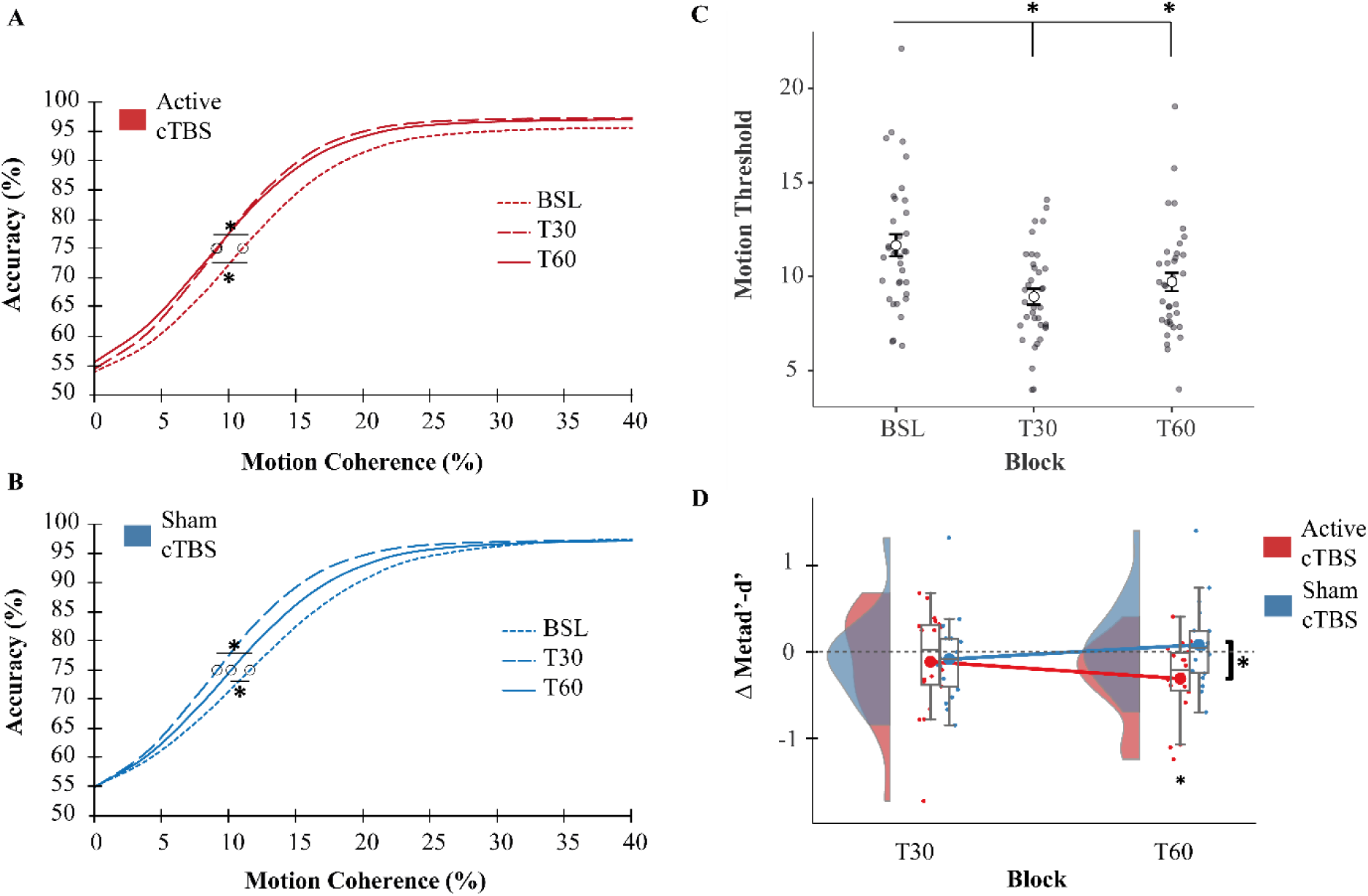
(**A,B**) Fitted data modeled on the logistic function to obtain the perceptual thresholds. The groups’ psychometric functions are separately plotted depending on the condition (in red shades Active_cTBS); in blue shades Sham_cTBS) and as a function of the testing Block (BSL, T30 and T60 Blocks). Circle dots depict the inflection points (IP) coincident with the percentage of coherent motion where the logistic function had a value of 75% of correct responses. **(C)** Raw motion sensitivity thresholds showing the Main effect of Block, values reported are collapsed mean and SEM with individual estimates for both groups, significant post-hoc comparisons are flagged with asterisks. **(D)** Changes in metacognitive efficiency following ccPAS and cTBS. Half-density and Box-plots with individual data points represent delta metad’-d’ at T30 and T60 for active (red) and sham (blue) condition of cTBS; Asterisks show significant one-sample t-test (below single condition) and post-hoc comparison (bars between conditions) respectively.

Importantly, and core to our hypothesis and experimental manipulations, average confidence levels (i.e. metacognitive bias) appeared to be affected by the interaction between cTBS with ccPAS (F(1,34)=4.81; p=.035; ηp^2^=.12; Fig.1C). Thus, the comparison between Block_T30_ and Block_T60_ in the cTBS_Active_ group revealed a significant decrease of average confidence (p=.016), whereas no changes in metacognitive bias are observed in the cTBS_Sham_ between Block_T30_ and Block_T60_ (p=.57). In detail, while we confirm changes in metacognitive bias relative to baseline (i.e., overconfidence) in the Block_T30_ following ccPAS (cTBS_Active_= 0.28±0.1, p<.01; cTBS_Sham_= 0.35±0.1, p<.01), this is selectively impacted in the Block_T60_ when IPS/LIP activity is inhibited (mean= 0.12±0.1, p=.12) – the subjects became underconfident of their performance - as opposed to sham cTBS (mean= 0.38±0.1, p<.01). Furthermore, other metacognitive abilities were partially modulated by the IPS/LIP manipulation. Specifically, for the metacognitive efficiency – i.e., the matching between performance accuracy and confidence level (see *Methods*) - we found a significant interaction Block x cTBS (F(1,34) = 6.88; p =.01; np =.17) (Fig 2D) without any other significant effect (Fs<1.79; Ps>.19). Posthoc revealed how participants that underwent the inhibitory cTBS (i.e., cTBS_Active_) showed a reduced efficiency in metacognition relative to sham stimulation of IPS/LIP (i.e., cTBS_Sham_) at Block_T60_ (p=.041). However, we did not find any significant modulation of efficiency from Block_T30_ to Block_T60_ in any of the groups (ps>0.09). Nonetheless, the examination of average efficiency shows that only following the administration of cTBS, a significant impairment could be observed (cTBS_Active_, T60= −0.32±0.1, p=.017), and this could not be noticed in any other Block (ps>.59, Fig. 2D). This confirmed IPS/LIP role in shaping second-order performance [15]. Interestingly, given that post-ccPAS metacognitive bias (i.e. confidence level) was generally increased compared to baseline levels (Ps < .015, Fig.1C), except following the inhibitory cTBS (mean= 0.12±0.1, p=.12), it suggests that reduced metacognitive efficiency in Block_T60_ for cTBS_Active_ might be related to a drop in confidence levels (i.e., underconfidence) despite the increased sensory accuracy [53]. Lastly, the ANOVA on metacognitive sensitivity – i.e., the precision of confidence attribution given the available sensory information (meta_d’, see *Methods*)-showed no relevant main effect or interactions (Fs<1.71; Ps>.19). This result further supports the findings of our original study [15] showing no modulation of metacognitive sensitivity following the induction of plasticity in the network encompassing IPS/LIP.

## Discussion

Our results provide direct evidence of the functional dissociation between the systems underpinning sensory and metacognitive decisions, defining the roles of the V5/MT+-to-V1/V2 pathway and its relationship with IPS/LIP in motion perception. First, we replicate previous findings showing that enhancing V5/MT+-to-V1/V2 connectivity with ccPAS increases sensory performance and, secondarily, biases decision confidence [15,23]. Crucially, we provide novel causal evidence that these changes rely on the activity of distinct cortical substrates, which can be selectively targeted. While visual motion discrimination seems to be routed in the V5/MT+-V1/V2 pathway, independently from IPS/LIP activations, perceptual metacognition relies on the functional interaction between the V5/MT+-V1/V2 visual pathway and IPS/LIP.

Motion discrimination is strongly dependent on V5/MT+-V1/V2 network [54]. In contrast, inducing plasticity in IPS/LIP does not influence sensory discrimination, confirming the limited involvement of posterior parietal cortices (PPC) in perceptual sensitivity [13,15,55]. Remarkably, the increase of metacognitive bias associated with the perceptual gain induced by ccPAS_V5/MT+--V1/V2_ – firstly reported in our previous study, and later replicated by others [15,23] – is critically affected by IPS/LIP disruption. The IPS/LIP area is shown to be involved in the conscious elaboration of sensory information [56–59]. Accordingly, our results demonstrate that perturbing IPS activity leads to a clear impairment of subjective confidence in perceptual judgments (i.e., metacognitive bias) and it also influences the efficiency of metacognition. These results corroborate the relevant role of posterior cortices in the wider network involved in metacognitive evaluation [60], and add to the increasing neural evidence linking confidence signals to parietal activation [30,57,61–65].

In line with our hypothesis, changes in metacognition observed after strengthening the V5/MT+-V1/V2 connections with ccPAS are the secondary result of the enhanced motion elaboration, which, in turn, alters the readout of sensory information in IPS/LIP. This confirms that ccPAS is capable of inducing functional changes in key nodes connected with the manipulated network [23,24]. The altered metacognition that accompanies the visual improvement is likely attributable to the anatomo-functional influence of V5/MT+ area over IPS/LIP [66–68]– an idea not at odds with previous interpretations [27] relating the effect of MT+ microstimulation to confidence changes [26]. Such phenomenon might also be due to an increased functional influence of early visual areas directly targeting PPC [69–71], as previously suggested [23]. Since ccPAS was designed to strengthen connections reentering in V1/V2, it likely enhanced these early regions as convergent hubs [19,72,73], thereby amplifying their impact on interconnected regions such as IPS/LIP.

Relative to our previous report [15], we now demonstrate that direct IPS/LIP inhibition via cTBS reduces both metacognitive bias and metacognitive efficiency, while leaving metacognitive sensitivity unaltered. The absence of metacognitive sensitivity modulation is consistent with the selectivity of the ccPAS effects that have been demonstrated in the original study, in which strengthening IPS/LIP-V1/V2 pathway increased metacognitive bias and efficiency [15]. Here, the decreased efficiency – that represents an index of the extent by which an observer is aware of his performance – found after the active inhibition of IPS/LIP explains the participants’ drop in subjective confidence (i.e., metacognitive bias) despite the augmented motion perception due to ccPAS_V5/MT+-to-V1/V2_. That is, the negative scores in efficiency are the likely consequence of a level of confidence not calibrated to the heightened sensory discrimination, which suggests how these decision-making abilities are independently processed and further supports the dissociation between the functional roles of the targeted networks.

On the other hand, our results indicate that the impact of cTBS over IPS/LIP on metacognitive bias is more robust than its effect on metacognitive efficiency. This is highlighted by a significant reduction of participants’ confidence from ccPAS administration (Block_T30_) to cTBS application (Block_T60_), whereas metacognitive efficiency is found to be lowered by cTBS only when compared to baseline performance (see *Results*). It is therefore possible that, to consistently influence higher metacognitive functions, which plausibly rely on broader network interactions, stimulation approaches such as ccPAS might be more suitable [15], as they are specifically designed to target functional connectivity [18]. Accordingly, higher metacognitive ability (i.e., efficiency) during perceptual tasks has been shown to depend on the integrity of the superior longitudinal fasciculus (SLF, [74]), which runs from occipital and parietal lobes to ipsilateral frontal cortices [75]. This highlights the critical role of network communication in visual decision-making as also suggested by evidence of higher functional connectivity between frontal and visual regions during metacognitive elaboration [62]. Thus, NIBS techniques that primarily target a single area to induce local plasticity, such as theta-burst protocols [40], may be less effective in causing sustained behavioural alterations in higher metacognitive ability when targeting posterior cortices (e.g.,[76]).

Our findings also suggest that distinct functional nodes (and pathways) within the metacognitive network may subserve different post-decisional processes [77] – in keeping with hierarchical models of metacognition [7,78]. This could explain the different modulation observed across distinct metacognitive indices in the present study. However, to fully capture the dynamic flow of information underlying each metacognitive facet – and to characterise how neurostimulation leads to specific changes in these – future studies should combine advanced NIBS protocol (e.g., ccPAS [23]) with imaging techniques (e.g., EEG). Further research is also needed to isolate the specific functional role of the pathway connecting V5/MT+ with IPS/LIP in human perceptual decision-making, in order to assess the behavioural effects of its modulation by means of associative stimulation. However, the anatomical proximity of these two cortical sites raises specific methodological challenges that may limit the feasibility of such ccPAS interventions.

In conclusion, our findings support a functional dissociation between neural systems underlying sensory and metacognitive processes [2,3,15,79], although such circuits appear intrinsically interconnected to support efficient decision-making [29,80,81]. Our work further elucidates the role of V5/MT+ and IPS/LIP in perceptual decision-making, pointing to independent levels of computation, with the parietal cortex integrating sensory signals in order to construct a sense of confidence [15]. These findings also imply that different decision-making abilities might develop in parallel, despite partially overlapping neural representations [29], thereby opening the way to potential NIBS applications aimed at selectively enhancing perceptual or metacognitive skills.

